# Brain scans from 21297 individuals reveal the genetic architecture of hippocampal subfield volumes

**DOI:** 10.1101/299578

**Authors:** Dennis van der Meer, Jaroslav Rokicki, Tobias Kaufmann, Aldo Córdova-Palomera, Torgeir Moberget, Dag Alnæs, Francesco Bettella, Oleksandr Frei, Nhat Trung Doan, Ingrid Agartz, Alessandro Bertolino, Janita Bralten, Christine L. Brandt, Jan K. Buitelaar, Srdjan Djurovic, Marjolein van Donkelaar, Erlend S. Dørum, Thomas Espeseth, Stephen V. Faraone, Guillén Fernández, Simon E. Fisher D.Phil., Barbara Franke, Beathe Haatveit, Catharina A. Hartman, Pieter J. Hoekstra, Asta K. Håberg, Erik G. Jönsson, Knut K. Kolskår, Cand. Psychol., Stephanie Le Hellard, Martina J. Lund, Astri J. Lundervold, Arvid Lundervold, Ingrid Melle, Jennifer Monereo Sánchez, Linn C. Norbom, Jan E. Nordvik, Lars Nyberg, Jaap Oosterlaan, Marco Papalino, Andreas Papassotiropoulos, Giulio Pergola, Dominique J.F. de Quervain, Geneviève Richard, Anne-Marthe Sanders, Pierluigi Selvaggi, Elena Shumskaya, Vidar M. Steen, Siren Tønnesen, Kristine M. Ulrichsen Cand.Psychol., Marcel P. Zwiers, Ole A. Andreassen Lars, Lars T. Westlye

## Abstract

The hippocampus is a heterogeneous structure, comprising histologically distinguishable subfields. These subfields are differentially involved in memory consolidation, spatial navigation and pattern separation, complex functions often impaired in individuals with brain disorders characterized by reduced hippocampal volume, including Alzheimer’s disease (AD) and schizophrenia. Given the structural and functional heterogeneity of the hippocampal formation, we sought to characterize the subfields’ genetic architecture. T1-weighted brain scans (n=21297, 16 cohorts) were processed with the hippocampal subfields algorithm in FreeSurfer v6.0. We ran a genome-wide association analysis on each subfield, covarying for total hippocampal volume. We further calculated the single nucleotide polymorphism (SNP)-based heritability of twelve subfields, as well as their genetic correlation with each other, with other structural brain features, and with AD and schizophrenia. All outcome measures were corrected for age, sex, and intracranial volume. We found 15 unique genome-wide significant loci across six subfields, of which eight had not been previously linked to the hippocampus. Top SNPs were mapped to genes associated with neuronal differentiation, locomotor behaviour, schizophrenia and AD. The volumes of all the subfields were estimated to be heritable (h^2^ from .14 to .27, all p< 1×10^-16^) and clustered together based on their genetic correlations compared to other structural brain features. There was also evidence of genetic overlap of subicular subfield volumes with schizophrenia. We conclude that hippocampal subfields have partly distinct genetic determinants associated with specific biological processes and traits. Taking into account this specificity may increase our understanding of hippocampal neurobiology and associated pathologies.

## Introduction

The hippocampus plays a key role in learning, memory and spatial navigation.^1^ It is known to be particularly vulnerable to pathological conditions and implicated in several major brain disorders, most notably schizophrenia^2,3^ and Alzheimer’s disease (AD).^4^

The breadth of findings regarding the role of the hippocampus in behaviour, and its non-specific association with a range of brain disorders, may result from the fact that it is a heterogeneous structure, consisting of cytoarchitecturally distinct subfields which subserve distinct functions.^5,6^ Lesion studies, as well as intrinsic connectivity patterns, support a dichotomy between an anterior section, attributed a role in anxiety-related behaviours, and more posterior regions, important for spatial processing and cognition.^7^ There is also a gradient of extrinsic connectivity to both cortical and subcortical regions across the longitudinal axis superimposed on the hippocampal intrinsic connectivity organization, illustrating the complexity of hippocampal biology.^8^ First episode schizophrenia has been most strongly associated with the *cornu ammonis* (CA)1 region and the subiculum in the anterior hippocampus^9,10^, though with longer illness duration more posterior regions also appear affected.^11^ AD is also thought to be primarily associated with volume reductions in CA1 and subiculum, with the dentate gyrus (DG) and CA3 relatively spared,^12,13^ though opposing findings have also been reported.^14^ Imaging genetics studies have firmly established that hippocampal volume is a highly polygenic trait. Given the differences in cytoarchitecture, connectivity patterns, and functions of the hippocampal subregions, it is likely that the volumes of the different subfields also have different genetic determinants. This is supported by gene expression studies documenting strict boundaries between subregions with respect to their transcriptional profiles.^15,16^ Genome-wide association studies (GWAS) have identified and subsequently replicated several single nucleotide polymorphisms (SNPs) that are significantly associated with total hippocampal volume.^17–19^ These GWAS also showed that top SNPs show localized effects on specific subcortical brain regions,^18^ and specific hippocampal subfields,^19^ rather than global effects. A follow-up study failed to find evidence of genetic overlap between schizophrenia risk and total hippocampal volume,^20^ which may be partly explained by lack of anatomical specificity in the volumetric estimates, i.e. a more granular approach may be required.

Recently, Iglesias and colleagues constructed a new atlas of the hippocampus, based on ultra-high resolution MRI data using *ex vivo* samples.^5^ This atlas has been combined with an automated segmentation algorithm and released as part of the popular neuroimaging software suite FreeSurfer v6. An initial analysis of this new software in several large-scale neuroimaging datasets established that all subfields are highly heritable, and that eleven of the twelve subfields show strong test-retest and transplatform reliability.^21^

In this study, we explored the genetic architecture of each hippocampal subfield volume, as segmented by the algorithm released with FreeSurfer v6. We hypothesized that the greater specificity of these measures, compared to total hippocampal volume, should reduce noise and allow for more sensitive detection of SNPs in genome-wide association analyses. By covarying for total hippocampal volume, we expected to identify associations that are specific to one or some of the subfields, allowing for a more nuanced understanding of this heterogeneous structure. In addition, utilizing summary statistics from previous large-scale GWAS, we sought to characterize the genetic overlap amongst the volumes of the subfields, with other subcortical and cortical regions, and with a diagnosis of schizophrenia or AD.

## Materials and Methods

### Participants

We included data from 16 cohorts that had structural MRI and genome-wide genotypes available, listed in supplementary Table S1, amounting to a total sample size of 21297 individuals. The age range of the sample covered a large part of the lifespan (mean age 47.8 years, standard deviation (SD) 17.3, range 3.2 – 91.4), and 48.3% was male. Information on individual cohorts, including brain disorder diagnoses (n=1464, 6.9% of total), is given in the supplementary information (SI), together with figures illustrating the distributions of demographics and their relation with hippocampal volume. Each sample was collected with participants’ written informed consent and with approval by local Institutional Review Boards.

### MRI data processing

Extended information on MRI data handling, including processing and scan quality control (QC), is given in the SI. Briefly, T1-weighted MRI volumes were processed using the standard FreeSurfer recon-all stream (v.5.3, http://surfer.nmr.mgh.harvard.edu). Hippocampal subfield volume estimates were subsequently obtained by running the novel subfield segmentation algorithm that was released as part of FreeSurfer v6.0. This algorithm employs Bayesian inference in combination with a hippocampal atlas created through manual delineation of ultra-high resolution (0.13mm) images of *ex-vivo* hippocampal tissue.^5^ As a sensitivity analysis, we calculated the correlation between hippocampal subfield volume estimates obtained through the combination of FreeSurfer v5.3 and the novel v6.0 hippocampal segmentation algorithm with those obtained when FreeSurfer v6.0 was also used for the main segmentation, for fifty participants. These correlations ranged from .87 for the parasubiculum to .96 for the hippocampal tail, as more fully described in the SI.

### Genotyping and quality control

Genetic data were obtained at each site using commercially available genotyping platforms. We carried out phasing and imputation according to protocols in line with those applied by the ENIGMA consortium (http://enigma.ini.usc.edu), applying standard quality control settings, further described in the SI. We restricted our analyses to those with European ancestry, as determined through multidimensional scaling (MDS).

### Statistical analyses

We included all twelve subfields as outcome measures in the analyses, approximately from anterior to posterior: the parasubiculum, presubiculum, subiculum, *cornu ammonis* fields 1, 2/3, and 4 (henceforth referred to as CA1, CA3, and CA4), granule cell layer of the dentate gyrus (DG), hippocampus-amygdala-transition area (HATA), fimbria (a white matter structure), the molecular layer of the DG, hippocampal fissure, and the hippocampal tail. We defined total hippocampal volume as the sum of all structures, minus the hippocampal fissure. Since the volumetric and genetic correlations between both hemispheres were extremely high for all structures (nearly all>.90), we summed the estimates of both hemispheres together to reduce the number of analyses.

Prior to all analyses, we regressed out the effects of scanning sites, sex, brain disorder diagnosis, age, and ICV from each outcome measure. This was done through generalized additive model (GAM)-fitting in R (v2.4.0) on the total sample, estimating each outcome measure from these variables, and continuing with the residuals. We further removed all individuals ±4 SD from the mean on any of the hippocampal measures or ICV.

To correct for the multiple comparisons, we calculated the degree of independence between the volume estimates of the subfields plus total hippocampus, by generating a 13×13 correlation matrix based on the Pearson’s correlation between all pair-wise combinations. Based on the ratio of observed eigenvalue variance to its theoretical maximum, the estimated equivalent number of independent traits in our analyses was 7.70. We therefore divided the community standard^22^ nominal genome-wide significance threshold of 5×10^−8^ by this number, setting a threshold of 6.5×10^−9^.

### Genome-wide Complex Trait Analyses

We used Genome-wide Complex Trait Analysis (GCTA)^23^ to calculate SNP-based heritability of each of the GAM-residualized subfield volume estimates, as well as those of other subcortical regions and cerebral lobes produced by FreeSurfer’s subcortical^24^ and cortical segmentation^25^ streams. We additionally included the first four population components, calculated through MDS on the entire sample, as covariates to guard against ethnicity effects. GCTA employs a restricted maximum likelihood (REML) approach, fitting the effects of all common SNPs as random effects by a mixed linear model, to obtain an estimate of the proportion of phenotypic variance explained by genome-wide SNPs. We further applied bivariate REML to estimate the genetic correlation between all regions.^26^ Before analysis, we removed regions with high linkage disequilibrium (LD) from the genetic data and pruned it, using a sliding window approach with a window size of 50, a step size of 5 and an R^2^ of 0.2, leaving 133147 SNPs. The BIG cohort was not included in these analyses as we did not have the genetic data in-house; the sample size for these analyses was therefore n=18979.

### Genome-wide association analyses

We performed a meta-analyzed GWAS, using PLINK. We chose this method over a mega-analysis design to avoid assuming an equivalence of genetic effect sizes across the cohorts, which differed in terms of mean age and other aspects of their recruitment, with virtually no loss in statistical efficiency.^27^ We first carried out a GWAS within each sample for the GAM-residualized estimates of the volume of the total hippocampus and each of the 12 subfields. We included the first four population components, calculated through MDS within each sample, as covariates. For the subfields, we also included total hippocampal volume as a covariate. We subsequently combined the within-sample results using a fixed-effect, inverse variance-weighted, meta-analysis in PLINK.

The GWAS were carried out using all individuals (n=21297), including individuals with brain disorders (n=1464, 6.9%), to maximize power. We re-analysed the data excluding patients to verify that the detected effects were not driven by the inclusion of patients with brain disorders. The regression coefficients for SNPs with P<1*10-^5^ from the main GWAS analysis on total hippocampal volume, including patients, was highly correlated with the regression coefficients when excluding patients (Pearson’s r=0.87).

### Functional Annotation

We used the Functional Mapping and Annotation of Genome-Wide Association Studies (FUMA) platform for functional annotation of the GWAS results.^28^ Through the SNP2GENE function, significant SNPs were mapped to genes based on positional, expression quantitative trait loci, and chromatin interaction information from 18 biological data repositories and tools integrated into FUMA. The resulting set of prioritized genes was checked for overrepresentation in gene-sets of biological processes and GWAS catalogues with the GENE2FUNC function, using a hypergeometric test.

### Genetic overlap with Alzheimer’s disease and schizophrenia

We applied cross-trait LD score regression (LDSR)^29^ and conditional false discovery rate (FDR) analysis^30,31^ to investigate the genetic overlap of each of the subfields with schizophrenia and AD. For this, we used the summary statistics from the 2014 PGC2 schizophrenia GWAS^32^ and the 2013 IGAP AD GWAS.^33^ Each set of summary statistics underwent additional filtering, including the removal of all SNPs in the extended major histocompatibility complex (MHC) region (chr6:25–35 Mb) and the use of only Caucasian samples. We further minimized sample overlap by rerunning the hippocampal subfield GWAS without the ADNI cohorts for comparison with the AD GWAS, and by removing the TOP and HUBIN cohorts from the schizophrenia GWAS. For further explanation of these two techniques, see the SI.

## Results

### SNP-based heritability

The SNP-based heritability of each subfield’s volume estimate, as well as additional regions of interest, and the genetic correlations amongst them are shown in Figure 1. Heritability estimates for all subfields, on the diagonal, were highly significant (all p-values < 1×10^−16^), ranging from h^2^=.14 of the parasubiculum to h^2^=.27 for the hippocampal tail. Full test statistics of the heritability estimates for all regions are listed in Table S2. Based on their genetic correlations, most of the hippocampal subfields formed a cluster, which further included the amygdala. The cortical gray matter volumes of the cerebral lobes clustered together, as did the pallidum, caudate, and putamen, i.e. basal ganglia structures.

**Figure 1.**
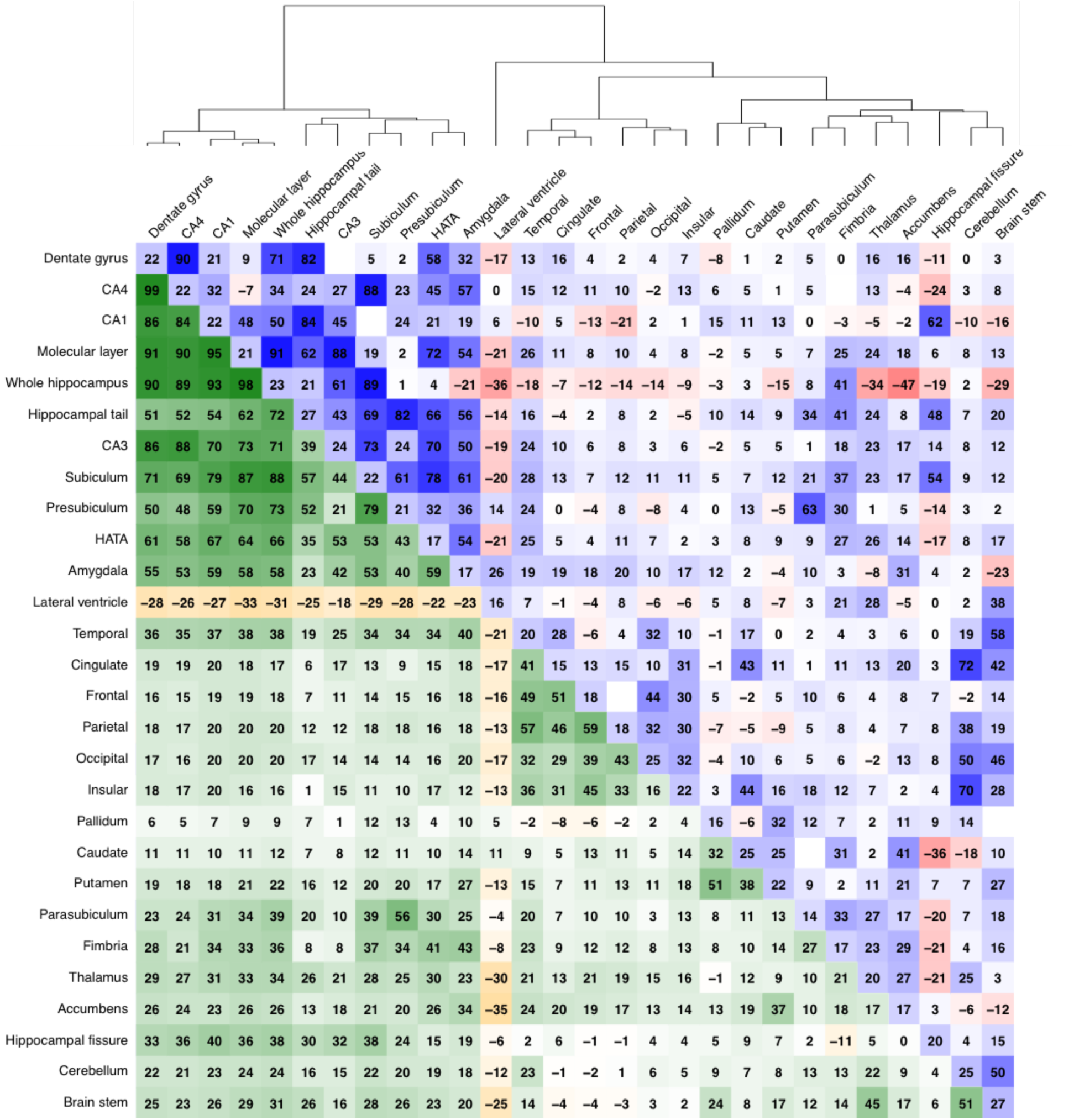
Correlation matrix of the volume estimates for the subfields as well as several other cortical and subcortical regions of interest and cerebral lobes. All correlations are multiplied by a factor 100. The volumetric correlations are shown in the lower triangle of the matrix (green-orange), the heritability estimates on the diagonal, and the genetic correlations in the upper triangle (blue-red). The order, indicated by the dendrogram on top, is determined by hierarchical clustering using Ward’s D2 method.

### Genome-wide association analyses

Our GWAS of whole hippocampal volume identified eight whole-genome significant loci. Of these, three loci have not been associated with the hippocampus before, namely those with lead SNP rs7630893 at chromosome 3 within the *TFDP2* gene, lead SNP rs2303611 within the *FAM175B* gene at chromosome 10, and rs1419859 at chromosome 12 upstream of *PARP11*.

The GWAS per subfield, corrected for whole hippocampal volume, identified a total of ten unique loci over six subfields. Of these ten, seven were not found for the GWAS on whole hippocampal volume. See Table 1 for information on each of the lead SNPs, per structure. Figure 2 provides an overview of the distribution of the p-values per top hit over the subfields, showing that while some have global effects, others are driven by specific subfields, most prominently the hippocampal tail. QQ plots and Manhattan plots for all subfields are shown in Figure S3. Forest plots indicated that all of the lead SNPs showed comparable effect sizes across the majority of cohorts, shown in Figure S4.

**Table 1.**
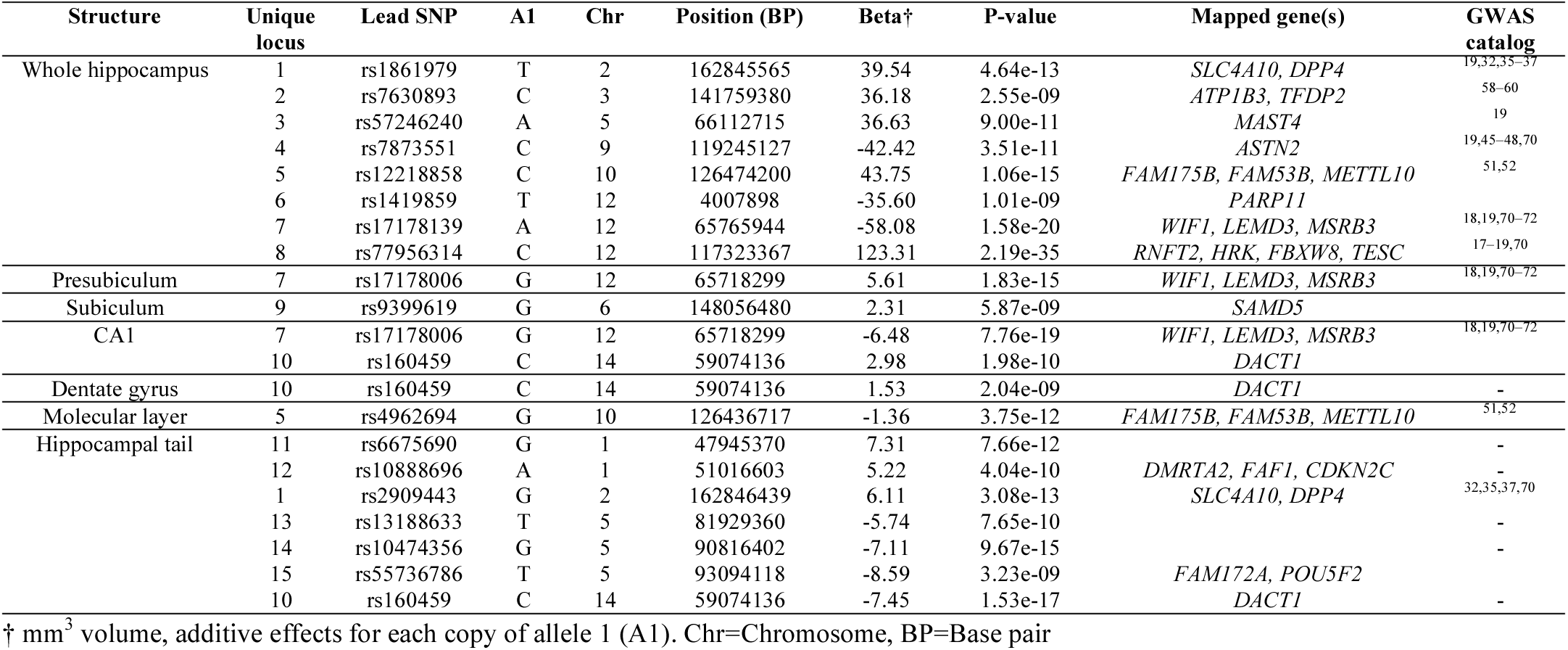
Whole genome significant loci for total hippocampal volume, as well as for the subfields while covarying for total hippocampal volume

**Figure 2.**
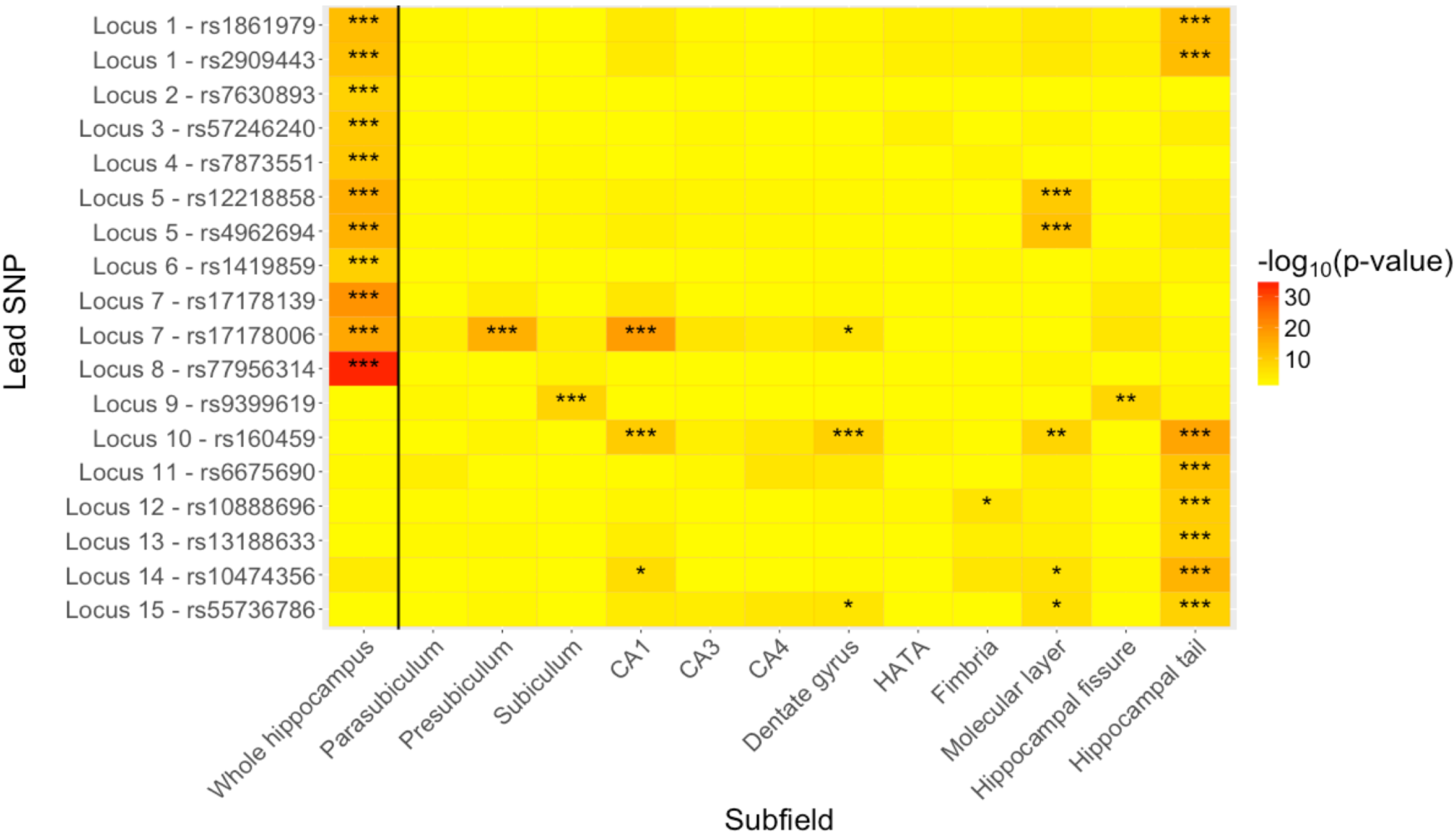
Heatmap based on the results from the genome-wide association analyses, showing the p-value for each of the lead SNPs reported in Table 2 (on the y-axis) per subfield (on the x-axis) volume. High −log10 p-values are shown in red, low values in yellow. Three stars in a field indicate the SNP reached whole-genome significance for that SNP (6.5×10^−9^), two stars nominal significance (5×10^−8^), and one star suggestive significance (1×10^−6^).

### Functional Annotation

The location of the genome-wide significant loci, in combination with the LD structure and known biological consequences of variation in these regions, led to the prioritization of 24 genes, listed in table 2 next to the loci that mapped onto them. Hypergeometric tests indicated that the lists of genes identified through the GWAS for both the volume of the whole hippocampus and the hippocampal tail were significantly enriched for genes associated with locomotive and exploratory behavior. Further, comparison to GWAS catalogs showed significant enrichment of AD-related genes for whole hippocampal volume, the hippocampal tail showed enrichment for schizophrenia-related genes, and the molecular layer was enriched for inflammatory bowel disease.

### Genetic overlap with AD and schizophrenia

Through LDSR, we found no significant evidence for genetic overlap of any of the hippocampal subfields with either disorder, as listed in Table S4. The conditional QQ plots did show enrichment as a function of association with schizophrenia for the presubiculum and subiculum, illustrated in Figure 3. This is not seen for other subfields, nor when conditioning on AD, see Figure S5. The subsequent conjunctional FDR analysis for these two subfields identified respectively five and four loci overlapping with schizophrenia, described in Table 2. Note that three out of nine hits have opposite direction of effects between subfield volume and schizophrenia, whereas the other six show the same direction of effects

**Figure 3.**
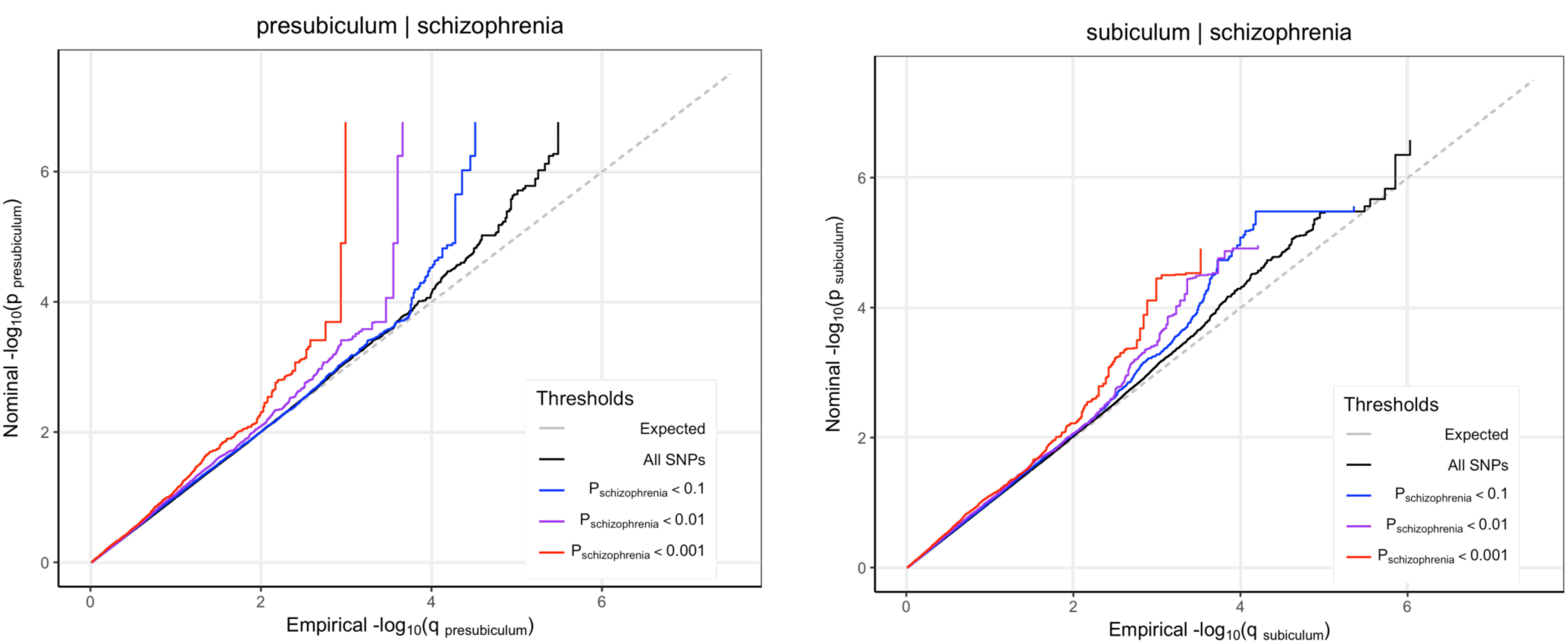
QQ-plots of the p-values from the presubiculum and subiculum genome-wide association studies (GWAS), conditioned on those from a schizophrenia GWAS. For both subfields, there is a clear upward deflection from the expected p-value distribution (in grey) that strengthens with increasing thresholds; the black line reflects the distribution of p-values from the subfields with no schizophrenia p-value threshold, blue shows the distribution of p-values remaining at a threshold of p<.1, purple those at a threshold of p<.01 and red those at p<.001.

## Discussion

The hippocampus complex comprises structurally and functionally distinct subfields with critical yet differential involvement in a range of behaviors and disorders. Using brain scans from 21297 individuals, we showed that differences in the cytoarchitecture of the subfields, providing the basis for their segmentation,^5^ are partly driven by differences in their genetic architecture. Further, greater specificity in the phenotypes under investigation allowed for the discovery of specific genetic variants. The elucidation of their genetic architecture, and identification of specific genetic variants, should be helpful in better understanding the biological functions of the individual subfields and their role in the development of common brain disorders.

The SNP-based heritability estimates we obtained, ranging from .1 to .3, were comparable to previous large-scale studies of the narrow-sense heritability of subcortical structures, when corrected for ICV.^20^ They also agree with findings from twin studies, showing that the larger subfields are the most heritable.^21^ We further found that the genetic correlations broadly mirror the volumetric correlations, and that the subfields cluster together with the amygdala. The strength of the correlations indicates that these structures share much of their genetic determinants, yet also confirm that they do indeed have specific, individual influences. Our estimates of genetic correlations with other structures corroborate findings from a twin study, that identified the same genetic clusters, with the hippocampus and amygdala clustering separately from respectively the cerebral lobes and basal ganglia structures.^34^

The genome-wide association analyses per subfield supported our reasoning that greater phenotypic specificity may aid genetic discoverability; we identified several genetic variants related to the volumes of subfields above and beyond total hippocampal volume. We found five out of six loci reported by a recent ENIGMA hippocampal GWAS, and the pattern of effects across the subfields also largely agree with their supplementary analyses of these top hits.^19^ This included a locus at chromosome 2 which maps onto the *SLC4A10* and *DPP4* genes, with our subfield analyses indicating this is driven by its effect on hippocampal tail volume. This locus has also been found in GWAS of educational attainment,^35^ cognitive ability,^36^ and schizophrenia.^32,37^ Further, inhibitors of DPP4 have been shown to improve recognition memory, lower oxidative stress and increase hippocampal neurogenesis in rodents.^38,39^ The well-known locus at chromosome 12, in the *MSRB3* gene, ^17,18,36^ on the other hand appears to be mostly driven by its effect on more anterior regions, being associated with the presubiculum and CA1. *MSRB3*, a gene involved in anti-oxidant reactions, has recently been shown to be particularly important for pyramidal neurons specifically in CA1, and to have lowered expression in the hippocampi of individuals with AD.^40^ The other locus on chromosome 12, linked to the *HRK* gene, appears to have a global effect, not being linked to any of the subfields after correction for whole hippocampal volume. *HRK* is a pro-apoptotic gene, associated with several forms of cancer^41^ and reported in one GWAS of AD age of onset.^42^ The two remaining replications, at chromosome 5 and 9, within the *MAST4* and *ASTN2* genes, also only appear for total hippocampal volume. *MAST4* codes for a microtubule protein part of the serine/threonine kinase family, with differential expression in frontotemporal dementia.^43^ *ASTN2* is thought to play a role in neuronal migration.^44^ It has been repeatedly associated with migraine,^45–48^ as well as schizophrenia^49^ and other neurodevelopmental disorders.^50^

The novel loci we identified may contribute to understanding the relation between certain peripheral diseases and cognitive dysfunction. The locus at chromosome 10, within the *FAM175B* gene, has been previously associated with cocaine dependence^51^ and bronchodilator responsiveness,^52^ as well as being reported in a recent GWAS of inflammatory bowel disease.^53^ Beyond whole hippocampal volume, it was found for the molecular layer of the dentate gyrus and the hippocampal tail, i.e. more posterior regions of the hippocampus. In rodents, lesions to the dorsal (corresponding to posterior in humans), but not ventral, hippocampus disrupt cocaine craving,^54,55^ and cocaine administration lowers neurogenesis in the dentate gyrus.^56^ Chronic intestinal inflammation has been associated with altered hippocampal neurogenesis, which has been theorized to explain the link between this disease and cognitive dysfunction.^57^ Another novel locus, at chromosome 3, lies within the *TFDP2* gene. This gene, with a function in cell proliferation, is well-known for its relation with kidney dysfunction.^58–60^ Chronic kidney dysfunction in turn is associated with cognitive impairment and hippocampal atrophy.^61^

Several genes were implicated through the GWAS on the subfields that were not identified for total hippocampal volume, illustrating the value of studying more specific phenotypes. Through the GWAS on the hippocampal tail, we found a locus at chromosome 1 with lead SNP rs4926555, within the *FAF1* gene. The protein product of this gene regulates neuronal cell survival and apoptosis,^62^ as well as glucocorticoid receptor-mediated transcription in hippocampal cells.^63^ The GWAS on the granule cell layer of the dentate gyrus and hippocampal tail further led to the identification of a novel locus at chromosome 14 with lead SNP rs160459, mapped to the *DACT1* gene. Knockout of *DACT1* has been shown to lead to decreased dendrite complexity in cultured hippocampal pyramidal neurons,^64^ and its expression has been linked to tumorigenesis suppression.^65^

Greater specificity in hippocampal segmentation also proved to be valuable for the investigation of genetic overlap with disorders. Through conditional FDR, we found signs of pleiotropy between schizophrenia and the subiculum and presubiculum, but not for other subfields. This is in line with studies showing that these anterior subfields are disproportionately affected in patients with first-episode schizophrenia.^9^ Such a distinction may indicate that the relation between the subicular regions and schizophrenia is more genetically driven, while the global reduction of hippocampal volume seen in later disease stages is relatively stronger influenced by environmental factors and the disease process. The subsequent conjunctional FDR analyses pinpointed some specific loci that overlapped, including *SLC39A8*, a gene well-known for its high pleiotropy,^66^ being linked to a range of traits besides schizophrenia, including cognitive functioning.^67^ These analyses also indicated that while some lead SNPs had opposing direction of effects on subfield volume versus schizophrenia, others had the same direction. These mixed directions of effects are indicative of a complex aetiology underlying the well-documented relationship between this disorder and hippocampal volume reductions. This will contribute to the null findings often reported on most global tests of genetic overlap,^20^ including our own LDSR analyses, as mixed directions of effects will cancel each other out. We further found no evidence of pleiotropy between AD and any subfield, despite the strong involvement of the hippocampus in this disorder. AD-related genes may influence hippocampal volume later in life through accumulation of effects over time, decreasing their effect size in the current study. Our pattern of findings once again illustrates the complexity of the genetic relationships between neuroimaging measures and disorders.

While our results are encouraging, future genetics studies may benefit from optimization of the subfield segmentation approaches. The segmentation algorithm employed here is based on an atlas created using histological and morphometric features.^5^ Gene expression studies of the hippocampus have indicated that there are numerous genetic domains with clearly demarcated borders that only partly overlap with this subfield division.^16^ We also found that the six subfields with significant loci were also the six largest subfields, i.e. subfield size appears positively correlated with discoverability of genetic variants. This pattern of findings likely partly reflects that the larger subfields are segmented with greater accuracy.^21^ Lastly, comparison of results with the literature is hindered by the differences in subfield definitions being used, and harmonization is needed^68^ to further improve discoverability.^69^

In conclusion, in addition to providing information on the localization of the effects on the hippocampus for previously identified genetic variants, we identified novel variants that influenced specific subfields. These variants were not previously associated with hippocampal volume, yet have known roles in neuronal differentiation and neurodevelopmental disorders. Together with the estimated genetic correlations, we have shown that hippocampal subfields have partly distinct genetic determinants, associated with specific biological processes and traits, thereby providing evidence that there is value in greater specificity of the brain phenotypes under investigation. Taking into account this specificity may aid in furthering our understanding of hippocampal neurobiology and associated functions and disorders.

## Acknowledgements

The research leading to these results has received funding from the European Union Seventh Framework Programme (FP7-PEOPLE-2013-COFUND) under grant agreement n° 609020 - Scientia Fellows; Research Council of Norway (#223273, 248778, 249711, 248980, 249795, 177458/V50); South East Norway Health Authority (#2013054, 2014097, 2015044, 2015073, 2016083, 2017112); The Kristian Gerhard Jebsen Stiftelsen, SKGJ_MED_008; and the European Community’s Seventh Framework Programme (FP7/2007–2013) under grant agreement #602450 (IMAGEMEND). This work further made use of data sharing from ADNI (funded by National Institutes of Health Grant U01 AG024904 and DOD ADNI Department of Defense award number W81XWH-12-2-0012), PING (National Institutes of Health Grant RC2DA029475), PNC (grant RC2MH089983 awarded to R. Gur and RC2MH089924 awarded to H. Hakonarson), and UKB (under project code 27412). Acknowledgments of funding sources for all cohorts participating in this study are listed in Table S3.

## Author contributions

D.v.d.M. and L.T.W. conceived the study; T.K., N.T.D., J.R. and L.T.W. pre-processed all data in FreeSurfer; N.T.D., M.J.L., C.L.B., L.B.N., L.T.W and T.K. QC’ed the data; D.v.d.M performed the main analysis with contributions from J.R., O.F., A.C.P., F.B., T.M., and L.T.W.; D.v.d.M and L.T.W. contributed to interpretation of the results. All remaining authors were involved in data collection at various sites as well as sample specific tasks. D.v.d.M. and L.T.W. wrote the first draft of the paper and all authors contributed to and approved of the final manuscript.

## Materials & Correspondence

The data incorporated in this work was gathered from various resources, see supplemental material. Material requests will need to be placed with individual PIs. D.v.d.M. and L.T.W. can provide additional detail upon correspondence.

